# Genealogies in Growing Solid Tumors

**DOI:** 10.1101/244160

**Authors:** Duke U. Rick Durrett

## Abstract

Over the past two decades, the theory of tumor evolution has largely focused on the selective sweeps model. According to this theory, tumors evolve by a succession of clonal expansions that are initiated by driver mutations that have a fitness advantage over the resident types. A 2015 study of colon cancer [44] has suggested an alternative theory of tumor evolution, the so-called Big Bang model, in which all of the necessary driver mutations are acquired before expansion began, and the evolutionary dynamics within the expanding population are predominantly neutral. In this paper, we will describe a simple mathematical model inspired by work of Hallatschek and Nelson [25] that makes quantitative predictions about spatial patterns of genetic variability. While this model has some success in matching observed patterns in two dimensions, it fails miserably in three dimensions. Despite this failure, we think the model analyzed here will be a useful first step in building an accurate model.

## 1 Introduction

In the 1950s Fisher and Holloman [20] and Nordling [41] found that within the age range 25–74, the logarithm of the cancer death rate increased in direct proportion to the logarithm of the age, with a slope of about 6 on a log-log plot. Nordling grouped all types of cancer together and considered only men, but the pattern persisted when Armitage and Doll [1] separated cancers by their type and considered men and women separately. Nordling [41] suggested that the slope of six on a log-log plot would be explained if a cancer cell was the end result of seven successive mutations.

In the studies cited above, the stages were unspecified events. That changed in 1971 with Knudson’s study of retinoblastoma [31]. Based on observations of 48 cases of retinoblastoma and published reports, he hypothesized that the disease is a cancer caused by two mutational events. He based this on the observation that the incidence time was exponential for the patients with multiple bilateral tumors, but looked like a gamma(2,λ) distribution for the less serious unilateral cases. He hypothesized that the more serious patients had a germline mutation in the underlying gene while the less serious patients needed to have mutations occur in both copies of the underlying gene. In current terminology the gene, later identified as *RB*1 is a tumor suppressor gene. Trouble begins when both copies are knocked out. For more on the development of these ideas see [32], which was written to mark the 30th anniversary of the original paper.

**Figure 1:**
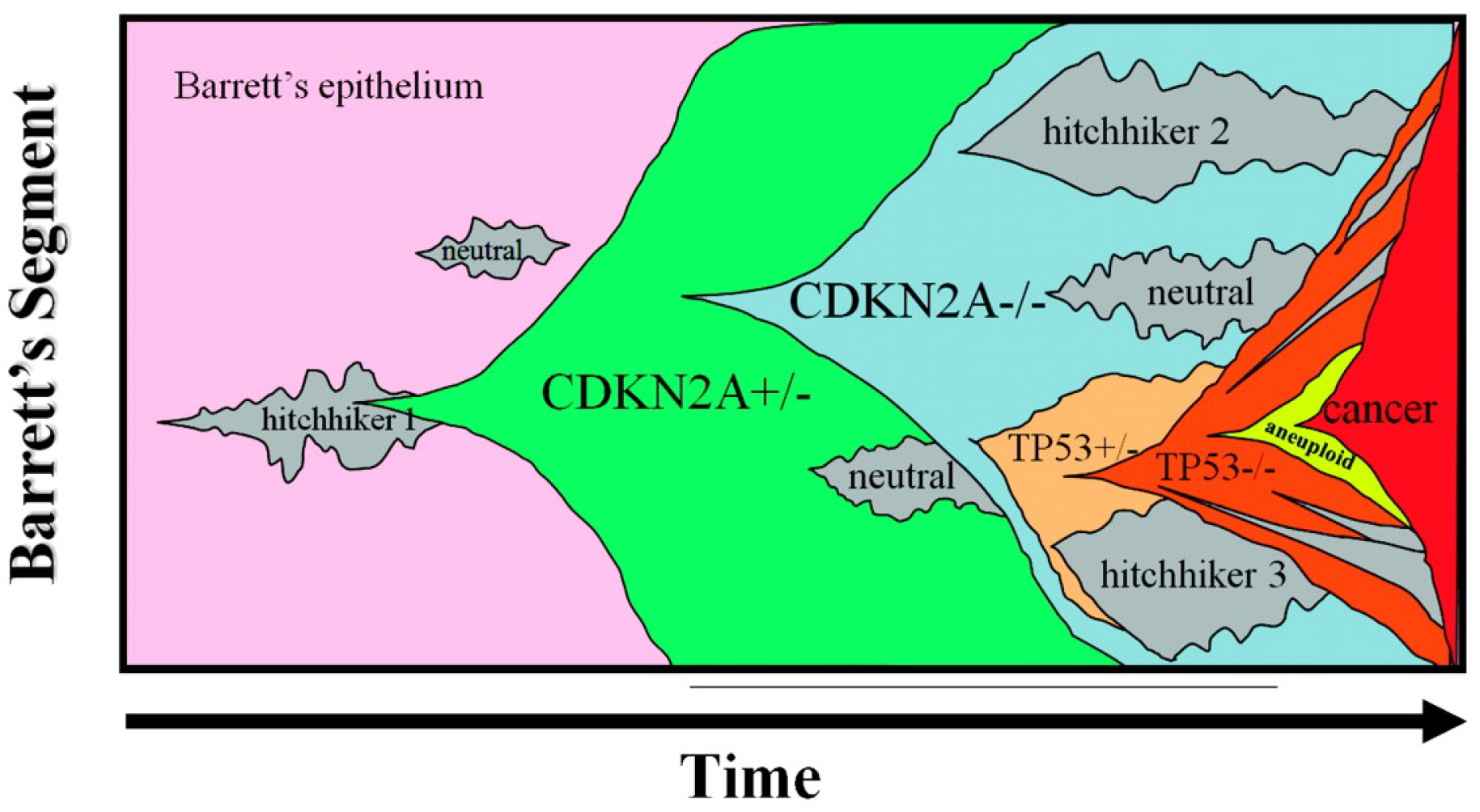
Clonal expansion in Barrett’s Esophagus. Figure is from [36].

In colon cancer the initiating event is thought to involve the inactivation of the tumor suppressor gene APC (adenomatous polyposis coli). In 1990, Fearon and Vogelstein [18] found a second piece of the puzzle when they noted that approximately 50% of colorectal carcinomas, and a similar percentage of adenomas greater than 1 cm have mutations in the RAS gene family, while only 10% of adenomas smaller than 1 cm have these mutations. In the modern terminology, the members of the RAS family are *oncogenes*. A mutation in a single copy is sufficient to allow progression. The analysis in [18] also suggested a role for TP53 (which produces the tumor protein p53) in the progression to cancer. The protein p53 has been described as “the guardian of the genome” because of its role in conserving stability by preventing genome mutations. Mutations in the gene TP53 have since been implicated in many cancers, see [21] and [48]. Combining these ideas leads to a four (or five) stage description for colon cancer that is described for example in the books of Vogelstein and Kinzel [47], and Frank [19].

In the multistage theory of carcinogenesis, it is thought that the sequence of “driver” mutations produces a series of selective sweeps. This theory has been confirmed by whole genome sequencing of cancer cells. Ding et al [10] have identified the clonal structure of eight relapsed acute myeloid leukemia (AML) patients. In one patient the founding clone 1 accounted for 12.74% of the tumor at the time of diagnosis. The additional mutations in clones 2 and 3 may have resulted in growth or survival advantages because they were 53.12% and 29.04% of the tumor respectively. Only 5.10% of the cells were in clone 4 indicating that it may have arisen last. However, the relapse evolved from clone 4 with the resultant clone 5 having 78 new somatic mutations compared to the sampling at day 170.

A mathematical theory has been developed for the clonal expansion model to make predictions about the level of intratumor heterogeniety [4, 15], the number of passenger (neutral) mutations in cancer cells [2, 46], and more practical questions such the evolution of resistance to treatmentt [26, 35, 45], the effectiveness of combination therapy [3, 29, 38], and the potential effectiveness of screening to reduce ovarian cancer [9]. See [13] for an introduction to the mathematics underlying many of these applications.

**Figure 2:**
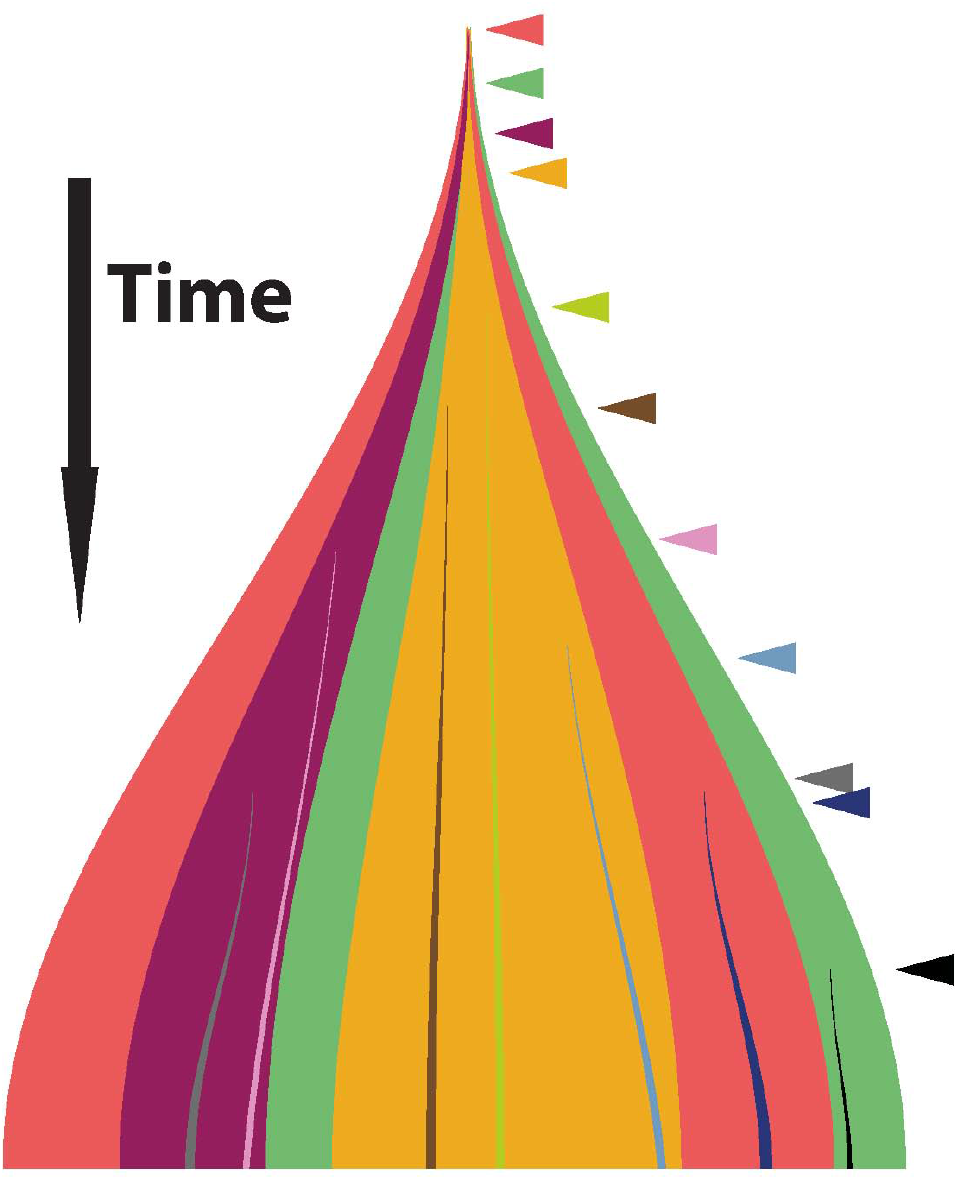
Cartoon picture of the Big Bang theory. Arrows mark the times of mutations. Colors indicate their spatial extent.

The picture of successive selective sweeps has been confirmed by comparing primary tumors and their metasases [50], and by regional sequencing of breast cancer [39], glioblastoma [43], and renal carcinoma [22]. Thus it was surprising when Sottoriva et al introduced and validated a ‘big Bang’ model in which all driver mutations were present at the time of tumor initiation. They collected genetic data of various types from 349 individual tumor glands were sampled from the opposite sides of 15 colorectal tumors and large adenomas. Data presented in Figure 3 of their paper shows that adenomas were characterized by mutations and copy number aberrations (CNA) that segregated between tumor sides. In contrast the majority of carcinomas exhibited the same private CNA in individual glands from different sides of the tumor. Follow up work of Ryser et al [42] found evidence of early abnormal cell movement in 8 of 15 invasive colorectal carcinomas (“born to be bad”) but not in four benign adenomas

**Figure 3:**
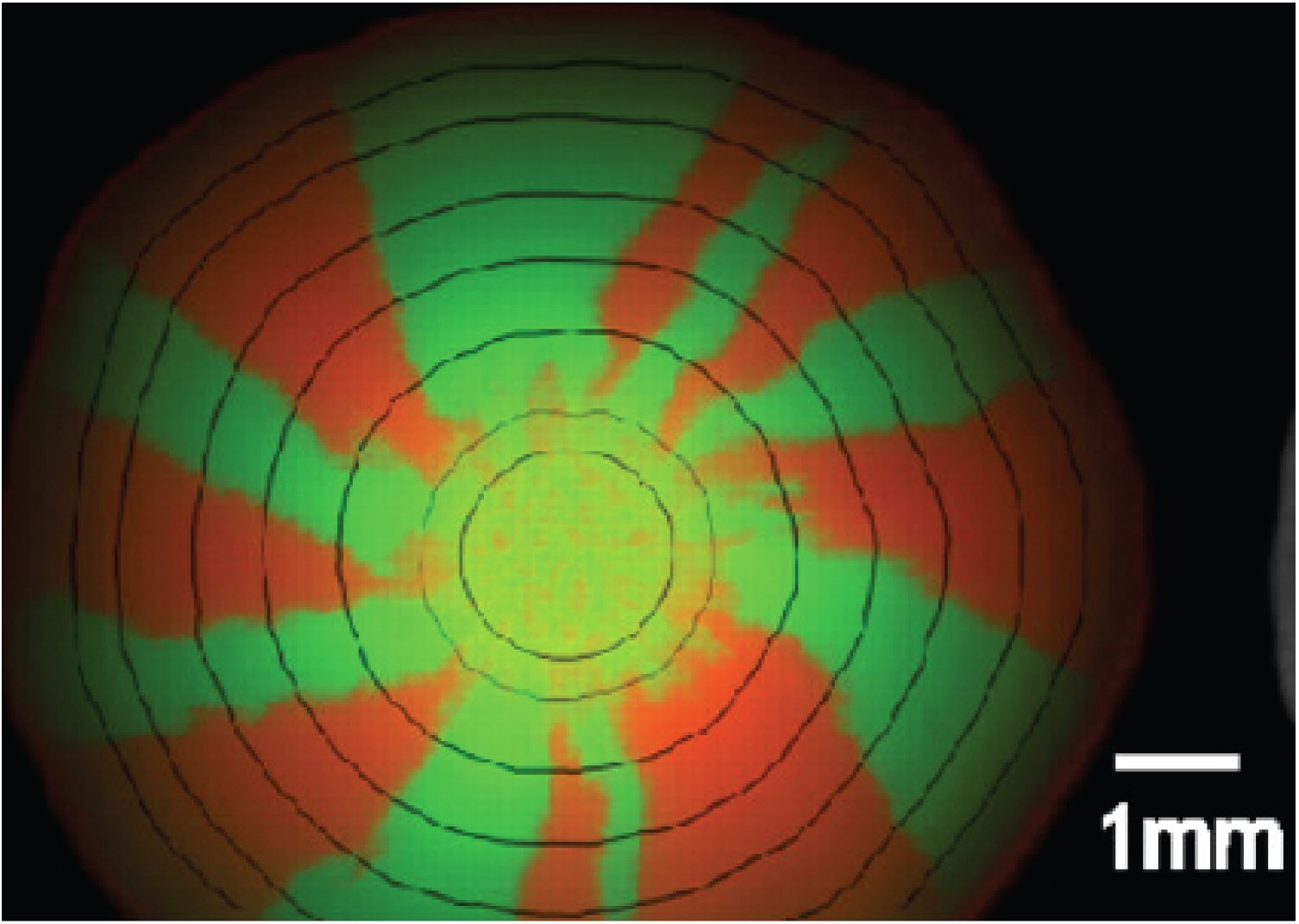
Yeast growth patterns.

The mutation patterns found by Sottoriva et al. [44] are similar to those found by Hallatschek et al [23] in a remarkable experimental paper. Two fluorescently labeled strains of *E. coli* were mixed and placed at the center of an agar plate containing a rich growth medium. The central region of the plate exhibits a dense speckled pattern reminiscent of the initial mixed population. From this ring toward the boundary of the colony, the population segregates into single colored sectors with boundaries that fluctuate. [25] and [30] have developed and analyzed models for the development of sectors in the system.

## 2 Biased Voter Model

A natural first step is to investigate genealogies in the biased voter model, which was introduced by Williams and Bjerknes [49] as a canceer model in 1972. In contrast to Durrett, Foo and Leder [16] there will be only two types of cells: 0 = wild type, 1 = cancer cells, and there are only neutral mutations, i.e., no type 2’s will be created, and there are no new mutations from type 0 to type 1.

In the biased voter model, 0’s give birth at rate 1, and 1’s at rate λ. In either case the new individual is sent to a randomly chosen nearest neighbor on ℤ*^d^*. Since we are concerned with solid tumors we will be primarily concerned with *d* = 2, 3. To key to analyzing the biased voter model is its duality with a branching coalescing random walk *ζ_t_*, see [6, 5]. To explain the duality we have to construct the process. For each *x ∈* ℤ*^d^* and nearest neighbor *y* we have a Poisson process 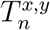 with rate 1/2*d*, and a Poisson process 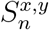 with rate (λ − 1)/2*d*. At times 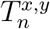 we put a *δ* at *x* and draw an arrow from *y* to *x*. The arrow reaches the the point *x* just above the *δ*. At times 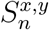 we draw an arrow from *y* to *x*. We think of *δ*’s as dams that stop the flow of fluid, and arrows that spread the fluid in the direction of the arrow. If we inject fluid at the bottom at the sites of *A* and allow the fluid to only flow up, being blocked by *δ*’s and flowing across arrows in the direction of their orientation the sites reached by fluid at time *t* are 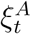. See Figure 4 for an example.

**Figure 4:**
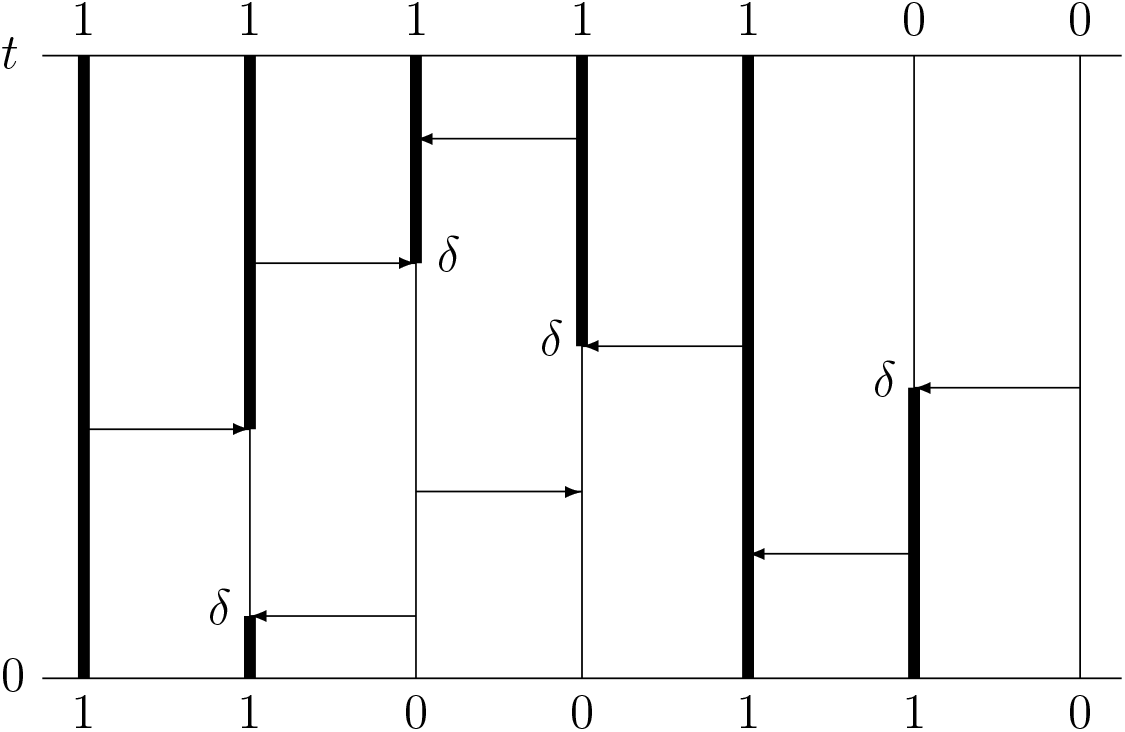
Construction of the biased voter model. Thicklines show the flow of fluid and indicate the (*x, t*) with ξ*_t_*(*x*) = 1.

To define the dual 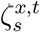, 0 ≤ *s* ≤ *t* we start at *x* at time *t*. Fluid flows down, is blocked by *δ*’s and moves across the arrows in the opposite direction to their orientation. From the definition it should be clear that *x* ∈ *ξ_t_* if and only if some site in 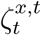 is occupied at time 0. Tat is,

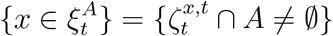

To define the dual starting from a set of sites *B* let 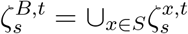. It is immediate that

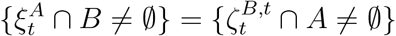

From the construction it is easy to see in the dual, particles jump to each nearest neighbor at rate 1/2*d*, give birth onto each nearest neighbor at rate (λ − 1)/2*d*. This system is called a coalescing branching random walk

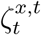 gives the set of potential ancestors of the individual at *x* at time *t*. To find the actual ancestor, we could follow the approach of the *ancestral selection graph* and starting from the values of *ξ*_0_ on 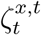 and work our way back up the graphical representations to see which arows produced births. Here we will follow the approach in Neuhauseer’s work on competing contact processes [40] and inductively define an ordering of the points in the dual 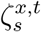 as we work backwards. The actual ancestor will be the first occupied site in the list. Looking at the example drawn in Figure 5 and working backwards from time *t*, the arrow from 0 to −1 will be an actual birth only if at that point −1 is vacant and 0 is occupied. To encode this we write the dual as −1, 0. The next two arrows as we work down is a voter arrow so the dual changes to −2, 0 and then to −2, 1. The arrow from −3 to −2 will be an actual birth only if at that point −2 is vacant and −3 is occupied, but if this is the case −3 will be the actual ancestor so the dual is now −2, −3, 1. Taking into account the last two arrows we see that the actual ancestor is the first occupied site on the list −1, −3, 1, 2.

**Figure 5:**
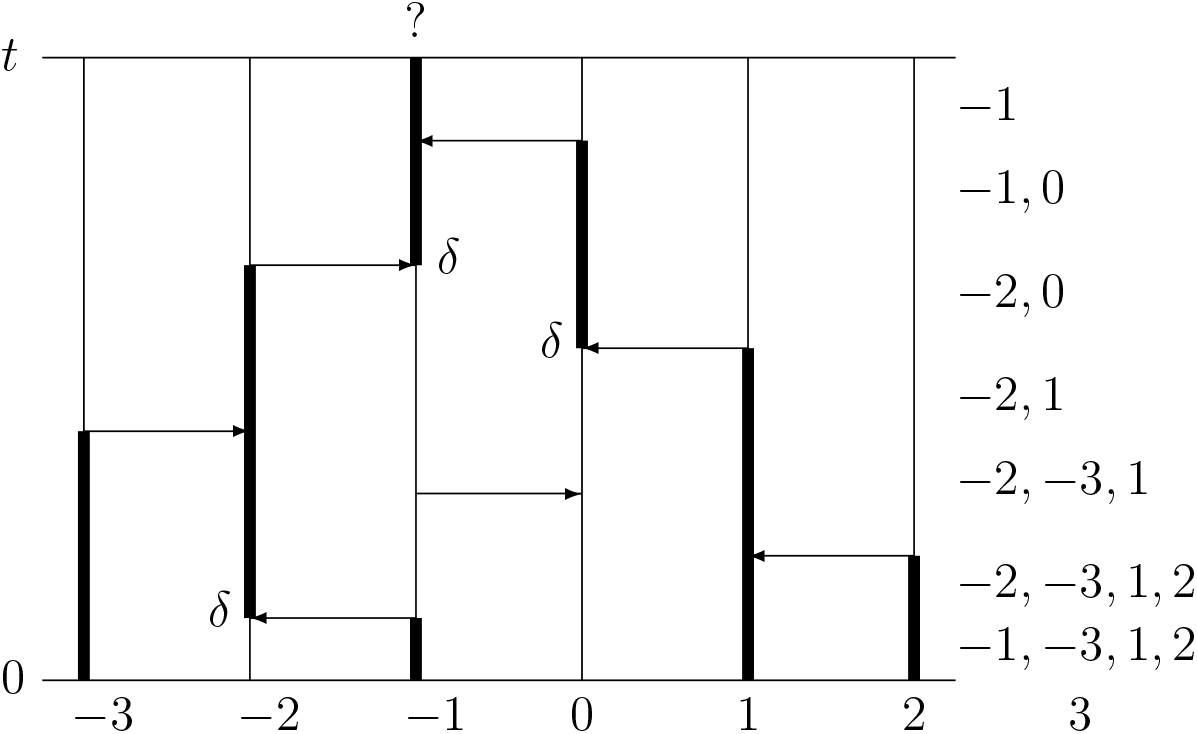
The dual 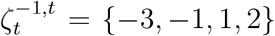 so −1 will be occupied at time *t* if one of these sites is.

Bramson and Griffeath [6, 5] showed that if we start with a single type 1 at the origin then when 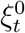 does not die out, it grows linearly and has an asymptotic shape *D*. That is, for any *∊* > 0, there is a *t_∊_* (which depends on the outcome ω) so that on {*T*_0_ = ∞} we have

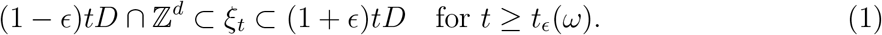

*D* is convex and has the same symmetries as those of ℤ*^d^* that leave the origin fixed, e.g., rotation by 90 degrees around an axis, or reflection through a hyperplane through the origin perpendicular to an axis.

If we select an *x* well away from the boundary at time *t* then initially all branching arrows connect two occupied sites. In this case they will not be part of the actual ancestral lineage, so it consists entirely of voter arrows. Individual trajectories will move like a mean zero random walk, and hence travel 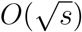 in time *s*. As Figure 6 shows, this means that the lineage will go almost straight down until it comes to the edge of the space time cone Γ = {(*x,t*): *x ∈ tD*}. At this point the lineage will begin to contain branching events that will take it back toward the origin. It is far from obvious what the path is doing when it is near the boundary but it must get to the origin since *x* ∈ *ξ_t_* implies 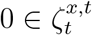.

**Figure 6:**
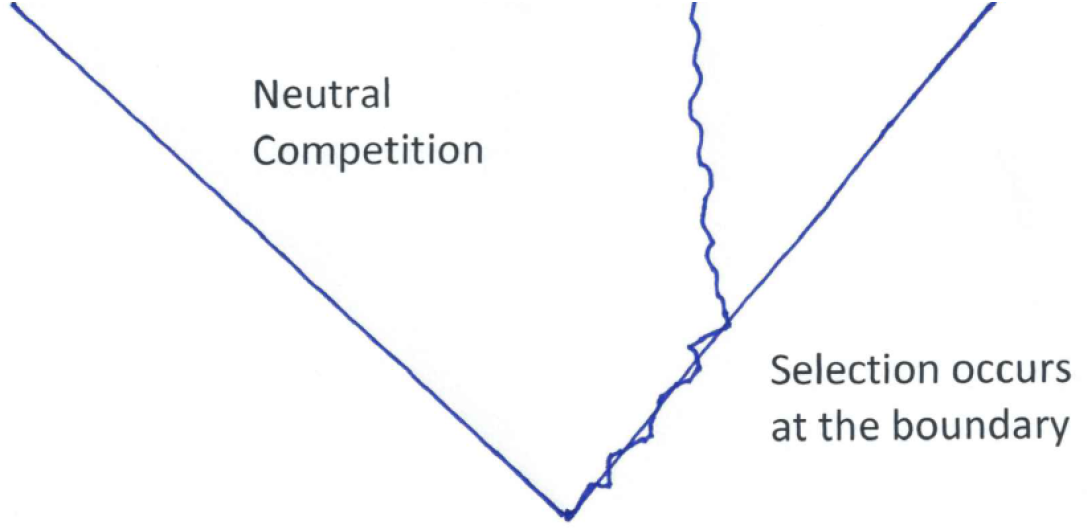
Behavior of the ancestral lineage of a cell in the biased voter model.

## 3 A simplified model

Hallatschek and Nelson [24] studied genealogies in a one dimensional system in which sites (demes) can carry up to *N* individuals. Here we will follow the formulation of Durrett and Fan [14]. There is one cell at each point of 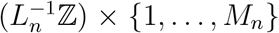, whose cell-type is either 1 or 0. The cells in deme 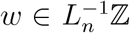 only interact with those in demes 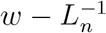 and 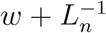 and interact equally with all of those points. Hence each cell *x* = (*w,i*) has 2*M_n_* neighbors. Type-0 cells reproduce at rate 2*M_n_r_n_*, type-1 cells at rate 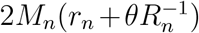. When reproduction occurs the offspring replaces a neighbor chosen uniformly at random. We define the approximate density by

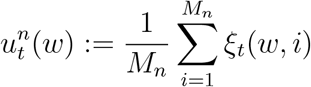
and linearly interpolate between demes to define 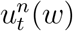 for all *w* ∈ ℝ.

let *C_b_*(ℝ) be the set of bounded continuous functions equipped with the metric

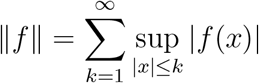
which induces the topology of uniform convergence on compact sets.

#### Theorem 1

Suppose that as n →∞, the initial condition 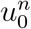 converges in C_b_(ℝ) to f_0_ and that:

(a) 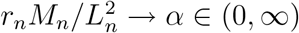
(b) 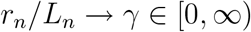
(c) 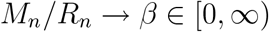
(d) 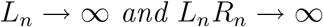

Then the approximate density process 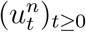 converges in distribution in D([0, ∞), C_b_(ℝ)), as n → ∞, to a continuous C_b_(ℝ) valued process (u_t_)_t≥0_ which is the weak solution to the stochastic partial differential equation (SPDE)

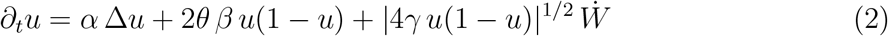
with initial condition u_0_ = f_0_. Here Ẇ is the space-time white noise on [0, ∞) × ℝ.

The absolute value is to make the coeficicent well defined when *u* < 0. When *θ >* 0 the solutions will stay nonnegative. For related results see the earlier work of Muller and Tribe [37] and Doering, Mueller, and Smereka [11] on limits of long range voter models To explain the terms in the limit

- 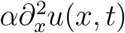 = diffusion generated by the voter model component of the dynamics
- 2*θβu*(1 − *u*) = increase in 1’s due to births from 1’s onto 0’s at rate *2M_n_θ/R_n_*
- 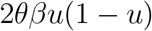 fluctuation due to random reproduction

Hallatschek and Nelson [24] viewed the system in a reference frame moving at rate *v* and made different parameter choices, so they ended up with the solution of the stochastic PDE

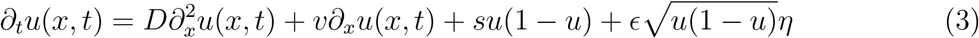
where *η* is space-time white noise, see their equation (2). The term *v∂_x_u*(*x, t*) arises due to the moving frame of reference.

Using arguments about the behavior of tracer particles placed into an expanding fluid the authors of [24] argue that the probability density *G*(*y, t/x, T*) that an individual at *x* at time *T* was descended from an ancestor that lived at time *y* at time *t* satisfies

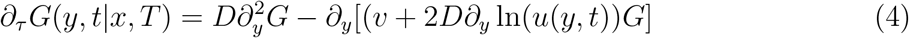

Recalling that a diffusion process with generator *L* = *a*(*x*)*fʺ*(*x*) + *b*(*x*)*fʹ*(*x*) has a transition probability that satisfies

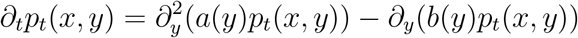
and comparing with (4), we see that in a fixed reference frame the coordinates of the ancestor will be a diffusion process with generator

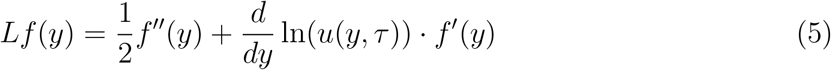
where *τ* = *T* − *t* since we are working backwards in time starting from *x* at time *T*. To see this is reasonable note that if the interface was as it is drawn in Figure 7 then in the flat part of the curve there would be no drift and the genealogy would be Brownian motion, in agreement with the heuristic picture in Figure 6. In the right side of the picture where *u* is decreasing, there will be a drift to the left, which will keep the lineage close to the boundary.

**Figure 7:**
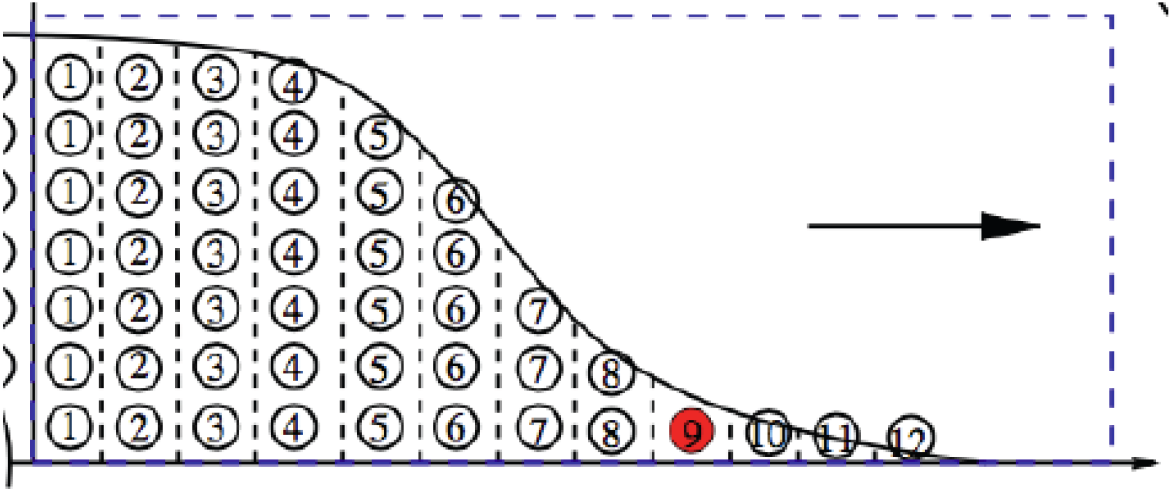
A one dimensional model with many individuals per site (or deme).

If *u* is the solution to (3), then it is not smooth enough for the drift (*d/dy*) ln[*u*(*y, t*)] in (4) to make sense mathematically. Even worse, in two dimensions there is no analogue of Theorem 3, since in dimensions *d* > 1 SPDE do not have function-valued solutions. To avoid these problems we could let 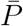 denote the probability measure for the biased voter model conditioned not to die out and switching to fixed reference frame replace the solution of (3) by

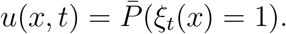

Since we do not know much about the right-hand side we will go one step further and let

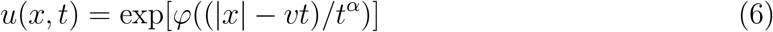
where *φ*(*z*) converges to 0 as *z* → −∞ and → −∞ as *z* → ∞. Here we have replaced the limiting shape in (1) by a ball and introduced *t^α^* as a measure of the fluctuations of the boundary. Thinking of the central limit theorem a natural choice is *α* = 1/2. However, physicists’ arguments [28] and simulations of the Eden model (which has births and no deaths) [51, 52, 34] suggest that in two dimensions the fluctuations of the boundary will be of order *t*^1/3^ rather than the usual *t*^1/2^. It is not known what the fluctuations should be in *d* > 2, so to accommodate a range of scalings we will allow *α* to vary and see how the behavior changes.

**Figure 8:**
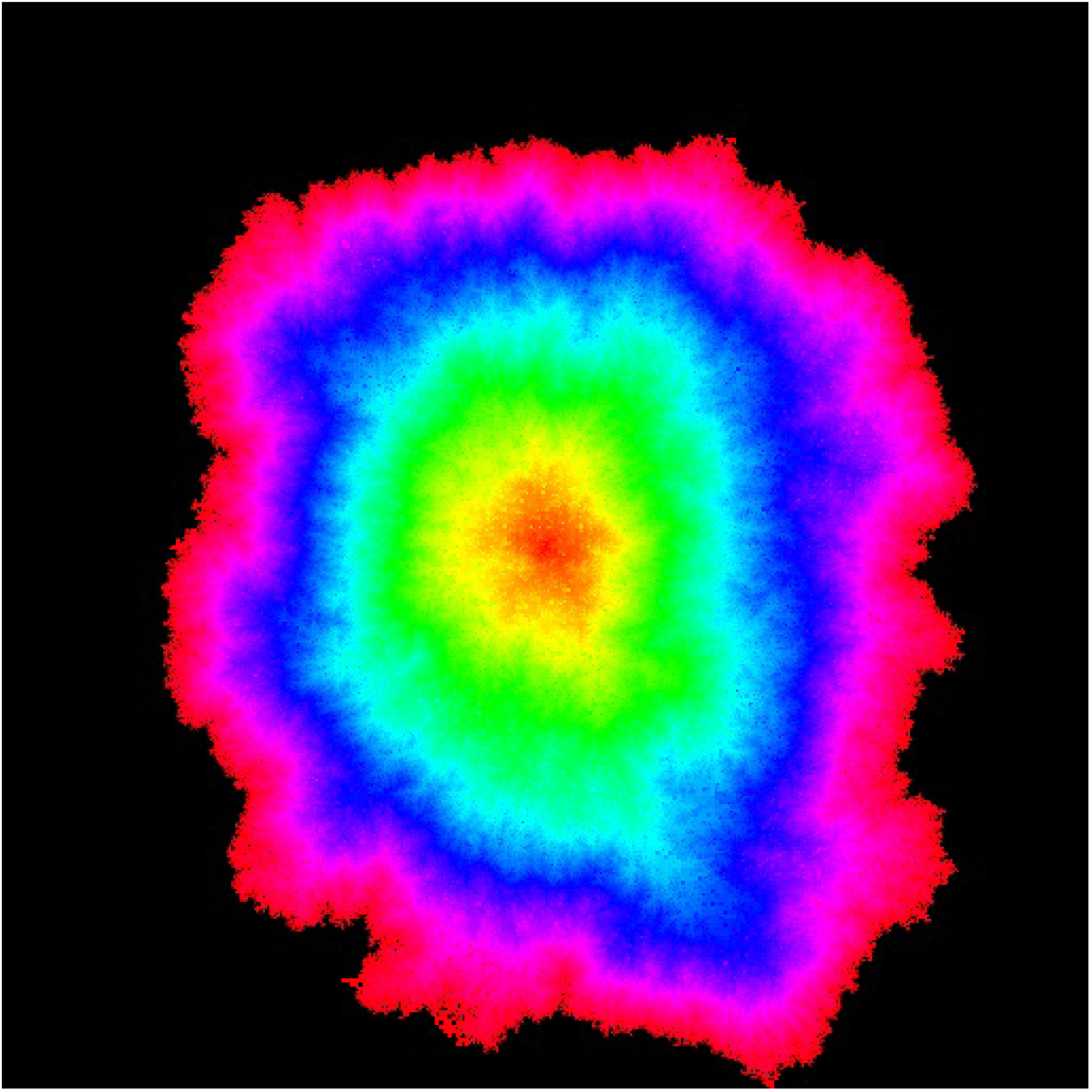
The Eden model has a very rough boundary.

Using the reasoning that led to (5), the process in *d* > 1 is:

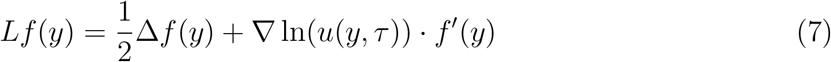
where again *τ* = *T* − *t*. Suppose, for the moment, that we are in two dimensions. Our next step is to change to polar coordinates. If we do this to Brownian motion (and add a superscript 0 to indicate that we are transforming Brownian motion) the result has

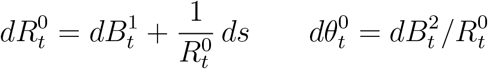
where the 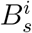 are independent Brownian motions. See [12] for this and other stochstic calculus facts that we use.

Since the drift in (7) is radial, in polar coordinates its angular part has

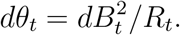

The radial component

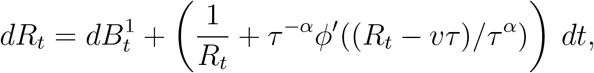
where *ψ* = *φ*ʹ/*φ*. Writing *U_t_* = *R_t_* − *v*(*T* − *t*) to return to the moving frame of reference, and dropping the first term which is small

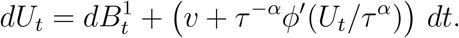

If we let *ψ*(*u*) = *vu* + *ϕ*(U/*τ^α^*) then we can write the last equation as

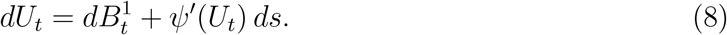

To understand the behavior of this process we note that its generator can be written as

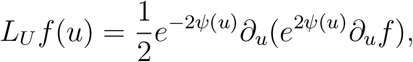
so if we let 〈*f, g*〉 = *∫ f*(*u*)*g*(*u*)*e*^2^*^ψ^*^(^*^u^*^)^ *du* then

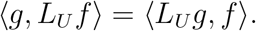

That is,

#### Lemma 1

L_U_ is self-adjoint with respect to e^2^ψ.

To have a concrete example we let

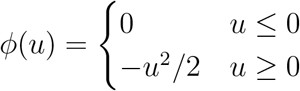
so that

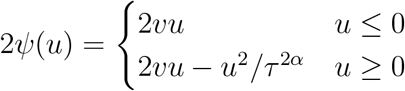
and the drift

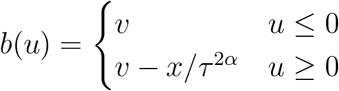

To make *e*^2*ψ*(*u*)^ a probability measure we have to normalize. To do this, we first suppose that the second half of the definition applies on the whole line and write

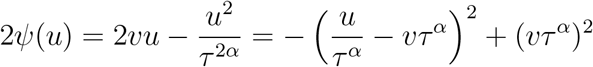
from this we see that

#### Lemma 2

When e^2ψ(u)^ is normalized to be a probability measure it is a normal with mean vτ^2α^ and variance τ^2α^/2.

From this, we see that it is legitimate to ignore the contribution from *u* ≤ 0. This result shows us that it is not sensible to take *α* > 1/2 since in that case the distance of the lineage from the tumor is much larger than the size of the tumor. The case *α* = 1/2 is also somewhat suspicious from this point of view since the displacement of the lineage from the boundary is of the same order as the diameter of the tumor. Later in Conclusion 2 we will see that when *α* = 1/2 all the coalescences occur near the beginning of tumor growth.

**Figure 9:**
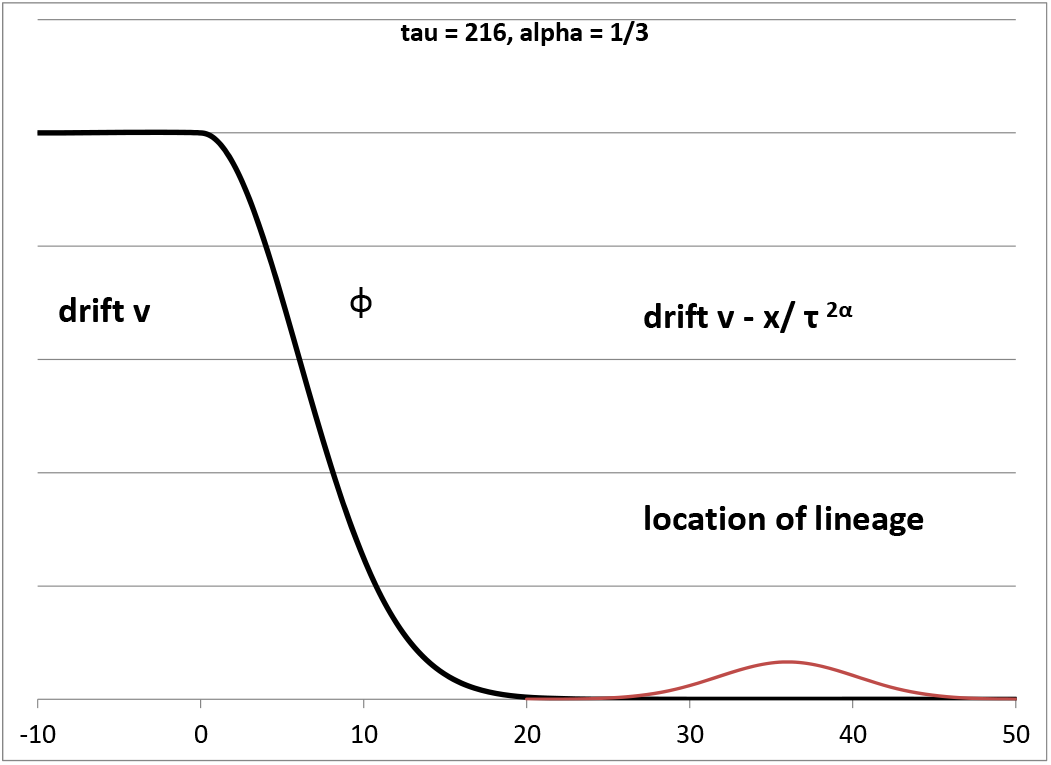
Picture of our concrete example indicating where the lineage is in equilibrium.

The reader should note that while (6) goes from 1 at *vτ* to ≈ 0 over a window of size *O*(*τ^α^*) the center of the stationary distribution is at *vτ*^2^*^α^*, which the point where the drift is 0. Because of this it is natural to look instead at 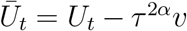. When we do this the drift is just −*ū*/τ^2^*^α^*, i.e., *Ū_t_* is an Ornstein-Uhlenbeck process.

## 4 Discretization of Radial Process

To have the possibility for lineages to coalesce, we will discretize *Ū_s_* and *θ_s_* to take place on *∊ℤ*. This is biologically natural since the tumor is composed of cells. To fix the units, we will think of space as measured in *cm*, and time as measured in years.

- Since cells have a diameter of roughly 10 *µm* = 10^−5^*m*, *∊* ≈ 10^−3^.
- Since tumors are several *cm* in diameter and take years to form the speed *v* ≈ 1. For simplicity we will set *v* = 1

The Ornstein-Uhlenbeck has unbounded drift, so a naive approach to discretization can lead to negative jump rates. To avoid this, we will instead use a continuous time birth and death process with stationary measure

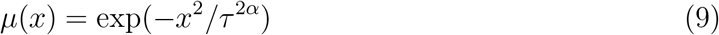
in which the rate of jumps from *x* to *y, q*(*x,y*), satisfies the detailed balance condition:

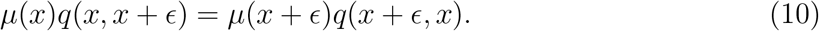

To have (10) we need

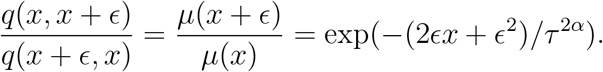

To do this we can take

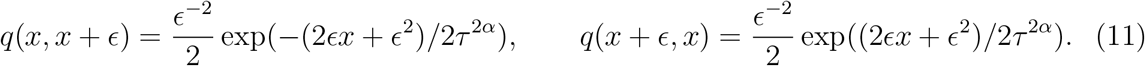

To see that discretized chain is reasonable note that replacing *x* by *x* − *∊* in the second formula this chain has infinitesimal mean

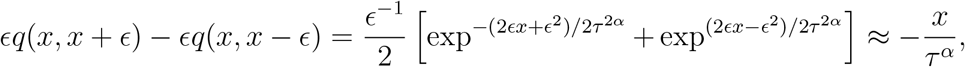
and the infinitesimal variance

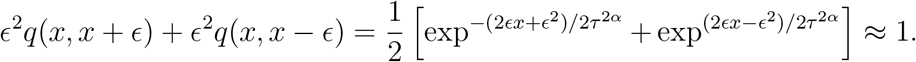

## 5 Angular Process

To discretize the angular process *θ_t_* when *R*_0_ = *R* we let 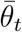 be a process that jumps *y* → *y* ± *∊/R* at rate 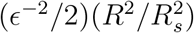 so that the infinitesimal variance is

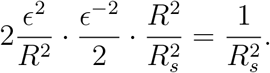

The angular variable is easy to understand. If we condition on *R_t_*, *t* ≥ 0 then

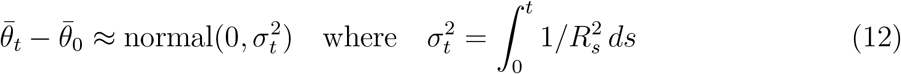

Using the fact that in the limit as *R* → ∞ *R_s_* ~ *T* − *s* = *R* − *s* we have

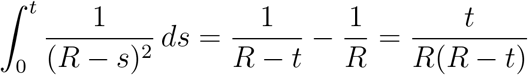

#### Remark 1

Note that for any δ > 0 this calculation is valid until t = R − R^δ^ because in that case σ^2^ = O(R^−δ^).

Consider now two points at distance *R* from the center whose angular parts differ by *θ*_0_ = *aR*^−1/2^. For the lineages can hit by time *t* we need

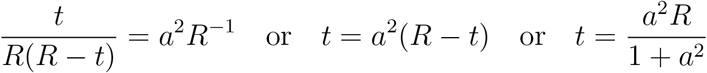

#### Conclusion 1

Tumor cells whose angles to the origin are separated by distance aR^−1/2^ will coalesce after a time with mean

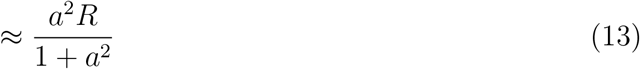

Due to the somewhat informal nature of our arguments, we will call them conclusions not theorems. To see what this says, note that when *a* = 3, *t* = 0.9R. If *R* = 5000 cells which is a tumor with of order 10^8^ cells and a radius 5 *cm*, 3R^1/2^ ≈ 210 cells. When *a* = 0.1 *t* ≈ 0.01R. In the concrete example 0.1*R*^1/2^ ≈ 7 cells

## 6 Coalescence of two lineages on the boundary

Consider now two independent genealogical processes 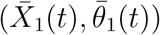 and 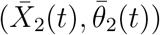. To have both coordinates of order *R* we will let 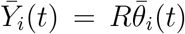 to get a process that jumps *y* → *y*±*∊* at rate 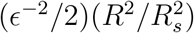. Let 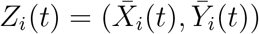. Let 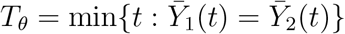, and let *T_c_* = min{*t* : *Z*_1_(*t*) = *Z*_2_(*t*)} be the coalescence time. Conclusion 1 gives a lower bound on the coalescence time by using *T_θ_* ≤ *T_c_*. We will now identify a situation in which *T_c_* occurs soon after *T_θ_*.

#### Conclusion 2

Suppose that

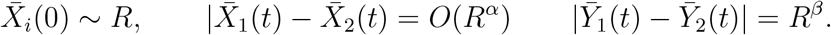

If *α* < *β* < 1/2.

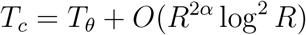

Since *T_θ_ = O*(*R*^2^*β*) *and α* < *β* this says *that T_c_ occurs soon after T_θ_*.

*Ideas that go into the proof*. In Section 8 we will show that the radial process comes to equilibrium in time *O*(*R^2α^*), and that in equilibrium 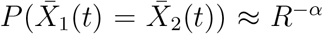. Because of this we can hope for coalescence when the amount of time the angular parts have been equal 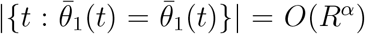. Since the probability of equality is *O*(*t*^−1/2^) this should take time *R*^2^*^α^*. However, due to the fact that two dimensional random walk is recurrent, we need time *O*(*R*^2^*^α^* log^2^ *R*). See Section 9 for details.

Note that the *Y* coordinates of lineages separated by *R^β^* with *β* < 1/2 will by conclusion 1 coalesce with high probability at a time that is *o*(*R*) so there will be solid patches of size *R^α^* by *R^β^* in polar coordinates in which most cells with the same ancestor.

## 7 Coalescence with interior lineages

#### Conclusion 3

Consider two lineages with

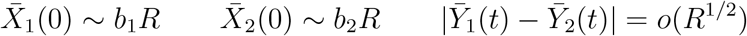
where b_1_ < b_2_. Then for any δ > 0 the probability the two lineages coalesce by time R − R^δ^ tends to 0.

*Ideas that go into the proof*. The closer lineage will do a two dimensional random walk until time (1 − *b*_1_)*R*. The first time that the two lineages are at the same distance from the center, the two angles will be separated by 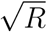, so by the argument in Section 9 they will need time at least *O*(*R* log^2^ *R*) to coalesce. The stated result comes from the fact that Remark 1 implies our calculations are only valid until time *R* − *R^δ^*.

Using results of Cox and Griffeath [8] we can study the case *b*_1_ = *b*_2_. The next result is a corollary of their formula (3.2). (The factor *θ*(1 − *θ*) comes from the fact that they consider the voter model starting from product measure with density *θ*.)

#### Conclusion 4

Consider two lineages with

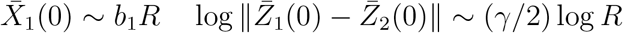
then the probability the two lineages coalesce by time t converges to 1 − γ.

The theorem says that the density of the cluster of sites whose genealogies coalesce with that of *Z*_1_(0) has a density that varies with the scale at which you examine it. Note that when *γ* = 1/2 the limit is 0. Using the hitting time results from Section 9 one can show that if 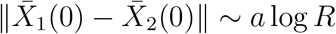 then the coalescence probability ~ *f* (*a*)/log *n*.

## 8 Equilibration of the centered radial process

When the initial separation is *O*(*R*^1/2^) the coalescence may or may not occur depending on the variability of the radial component. In this section, we will show that the time for the radial process seen from the moving reference frame to reach equilibrium is *O*(*τ*^2^*^α^*). As we have already shown in Lemma 2 the standard devition in equilibrium is *O*(*τ^α^*). Consider first times *t* ≪ *T*. In this case *τ* = *T* − *t* is constant and the radial process in the moving reference frame is

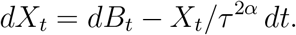

If we let *β* = *1/τ^2α^* to simplify notation, this SDE can be solved explicitly by

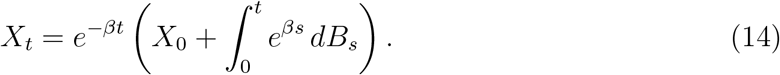

A proof can be found on page 180 of [12] but it is easily checked by formally differentiating the expression in (14)

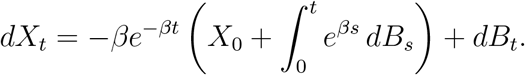

From this we see that when *X*_0_ = *x, X_t_* is normal with mean 0 and variance

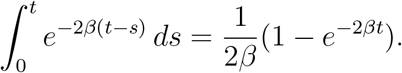

As *t* → ∞ this converges to normal(0,1/2*β*). In addition, we can see from the formula if *t* ≫ 1/*β* = *τ^2α^* we are close to equilibrium.

We need to derive a convergence to equilibrium for the discretized process. To lead up to this we will take another approach to the previous result. We start with the infinitesimal generator for the OU process

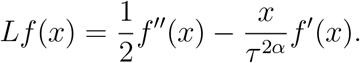

To look for an eigenfunction, we guess *f*_1_(*x*) = *x* which has

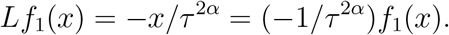

For the Ornstein-Uhlenbeck generator it is known that − 1/τ*^2α^* is the largest negative eigenvalue so the rate of convergence to equilibrium is *τ^2α^*.

To mimic this argument in the discrete setting note that what *µ* defined in (9) is not-malized to a be a probability measure it is approximately

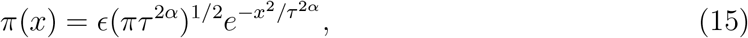
since summing over *x* gives a Riemann sum approximating the integral of a normal density. From the definition of the rates *q*(*x,y*) it satisfies the detailed balance condition. Thus the flow of probability across the edge (*x, x* + *∊*) in equilibrium is

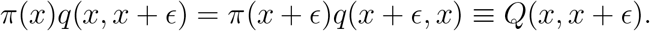

To bound the spectral gap we look at the Dirichlet form

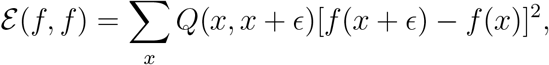
and note

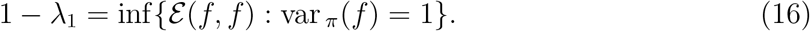

If we take *f*_1_(*x*) = *x* then var _π_(*f*) = τ^2/α^/2 while using (9) and (11)

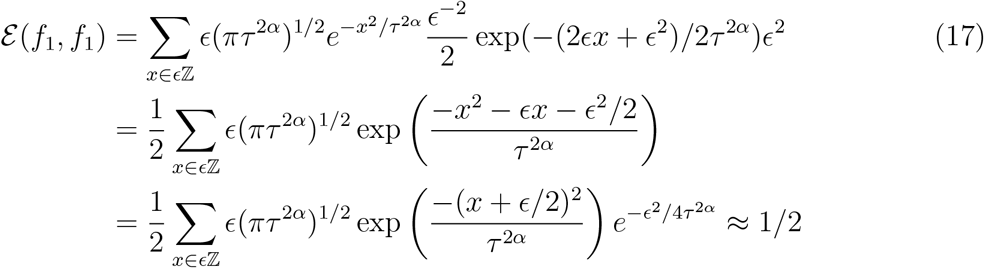
since when the last term is removed this is close to the integral of the density function for the normal with mean −∊/2 and variance *τ*^2^*^α^*/2. Using (16) now we conclude that in the discretized version the spectral gap is close to 1/*τ*^2^*^α^*, i.e., the distribution of 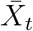 comes to equilibrium in time *τ*^2^*^α^* ≤ *T* since 2*α* < 1.

## 9 Estimating the Coalescence Time

We begin by explaining the method we will use, starting with the simpler problem of determining how long it takes for two independent continuous time random walks to hit on *ℤ*^2^. The methodology follows Section 3 of [53]. Let *W_t_* be the difference in the positions of the two walks and let *T*_0_ = min{*t* : *W_t_* = 0}. Breaking things down according to the value of *T*_0_ and writing the density function of *T*_0_ as *P_x_* (*T*_0_ = *s*)

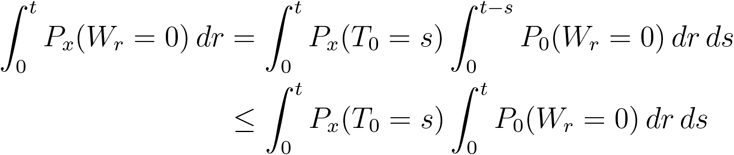
so we have

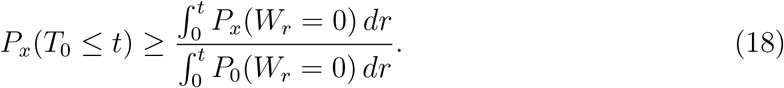

Note that in using this result we want a lower bound on the numerator and an upper bound on the denominator.

For a bound in the other direction we use

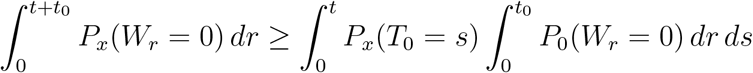
which gives

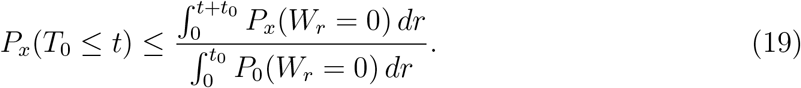

Note that in using this result we want an upper bound on the numerator and a lower bound on the denominator.

#### Proof of Conclusion 2

The assumption *α < β* implies that the centered radial component is in equilibrium at times *t* ≥ *R^2β^*. Suppose 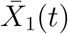 and 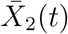 are independent and have the equilibrium distribution (15). Using the fact that *π*(*x*) = *π*(*−x*) on *∊ℤ* and is an approximation to normal(0,*τ*^2^*^α^*/2), we see that if *t* = *o*(*R*) then

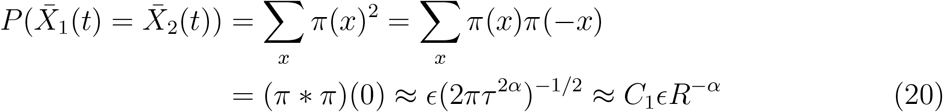
where * indicates convolution and *C*_1_ = (2*π*)^−1/2^ is a constant. Using this and the local central limit theorem we conclude that if *K* is large

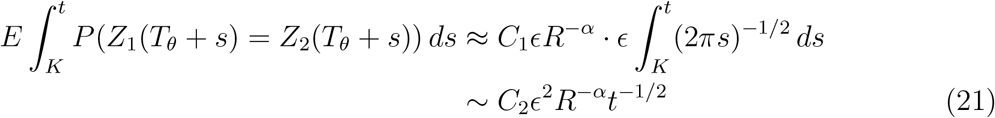
where *C_2_* = *C*_1_(2π)^−1/2^ · 2 = *π^−^*^1^

To get a lower bound on *P*(*T_θ_* ≤ *t*) using (18) we have to compute

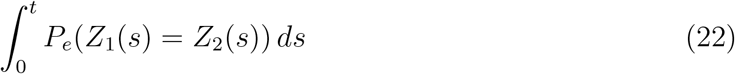
where *e* denotes the process starting from *Z*_1_(0) = *Z*_2_(0). The integrand is

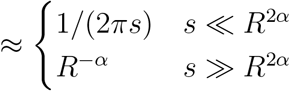
since in the first case the drift in the OU process is not significant and in the second the radial part is in equilibrium. In the gap between the two cases the formulas are almost the same, so the expression in (22) is

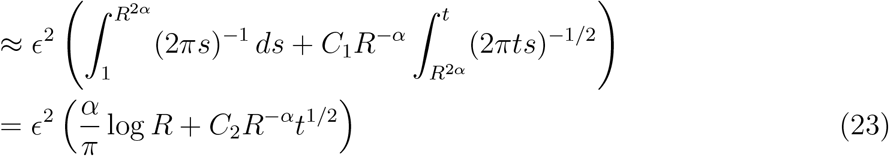

Combining the last result with (21) and using (18) we see that if *t* = *K R^2α^* log^2^ *R* then

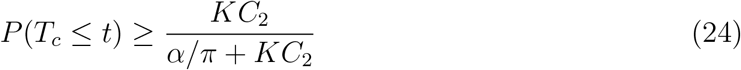
which is close to 1 if *K* is large. Since *T_c_* ≥ *T_θ_* we do not need a lower bound.

Conclusion 3 can be proved in almost the same way using (19).

## 10 Discussion

At this point we have achieved our goal of analyzing the behavior of the lineages in the simplified model and found that near the boundary the coalescence patterns are similar to those found in the tumor data. Unfortunately, some other properties do not agree with observed patterns

- The patches of cells that share the same genealogy with an interior lineage have diameter *O*(*R*^1/2^) are not dense enough to be detected by biopsy.
- The genealogies we follow are *τ*^2^*^α^* behind the front where the density of tumor cells is very small.

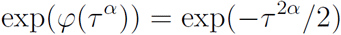 Taking *φ*(*u*) = *u*^γ^/γ with γ > 2 keeps genealogies closer but still the density at their location is low.
- In *d* = 3 if two cells are separated by log *R* at time *t* then, with high probability, they coalesce after time *R* − *R^δ^*, since three dimensional random walk is transient.

One of the problems with the current model is that there is considerable amount of turnover of cells in the interior of the tumor. This is not realistic since cell divisions in the interior are rare, and in some cases the cells will be hypoxic and die. We are currently investigating variants that only have growth at the frontier.

## Acknowledgements

This was partially supported by DMS 16164838 from the math biology program at NSF. The author would like to thank Jasmine Foo, Kevin Leder and Matthew Junge for useful comments. He would also like to thank David Basanta and Sandy Anderson for an invitation to visit the Integrated Mathematical Oncology at the Moffitt Cancer Center and an opporunity to give a talk about this work while it was still under development. Similar thanks go to Maury Bramson and the probability seminar at Minnesota.

